# Effect of RNA on morphology and dynamics of membraneless organelles

**DOI:** 10.1101/2021.04.16.440139

**Authors:** Srivastav Ranganathan, Eugene Shakhnovich

## Abstract

Membraneless organelles (MLOs) are spatiotemporally regulated structures that concentrate multi-valent proteins or RNA, often in response to stress. The proteins enriched within MLOs are often classified as high-valency “scaffolds” or low valency “clients”, with the former being associated with a phase-separation promoting role. In this study, we employ a minimal model for P-body components, with a defined protein-protein interaction network, to study their phase-separation at biologically realistic low protein concentrations. Without RNA multivalent proteins can assemble into solid-like clusters only in the regime of high concentration and stable interactions. RNA molecules promote cluster formation in an RNA-length dependent manner, even in the regime of weak interactions and low protein volume fraction. Our simulations reveal that long RNA chains act as super-scaffolds that stabilize large RNA-protein clusters by recruiting low-valency proteins within them while also ensuring functional “liquid-like” turnover of components. Our results suggest that RNA-mediated phase separation could be a plausible mechanism for spatiotemporally regulated phase-separation in the cell.

## Introduction

The cellular milieu is heterogeneous, and diverse intermolecular interactions can drive phase-separation of biological macromolecules leading to formation of spatially segregated membraneless compartments^1^. Membrane-less organelles (MLOs) are involved in several important cellular functions including ribosome synthesis and spindle formation during cell division^2^ A common feature of these membrane-free “organelles” is liquid-liquid phase separation (LLPS)^2,3^. One such class of biomacromolecular assemblies is that of RNA-protein (RNP) condensates which are present within both the nucleus and the cytoplasmic space ^4–8^. Examples of cytoplasmic RNP assemblies include P bodies, germ granules and stress granules (SGs) ^6,9,10^. Macromolecular assemblies formed by LLPS regulate gene expression by co-localising RNA and translation-related proteins within them^9,11,12^. Other functions attributed to RNP condensates include (i) sequestering mRNAs stalled during translation in response to stress ^9,13^, and (ii) segregating protein machinery involved in RNA processing and transport ^6,14^.

Experimental studies involving SGs identified the role of the granules, elucidated their constituents, and partially characterized structural dynamics of SGs^15,16^. While several experimental studies have shown the ability of RNA-binding proteins to assemble into droplets, these observations relate to overexpression of the protein of interest *in vivo*, or at large saturation concentrations *in vitro*^17–21^. This raises a critical question – is phase separation a relevant mechanism for assembly of MLOs in the cell at physiologically low protein concentrations? Are there other spatiotemporally regulated factors that are essential for their phase-separation at low abundances of phase separating proteins? In this context, the potential role of RNA as a potential LLPS promoter becomes critical. While RNA is found to be an integral component of several condensates (mRNAs in SGs and P-bodies, lncRNAs such as NEAT1 in paraspeckles) ^5,22–24^, the potential role of RNA molecules in regulating phase behavior and the dynamics of these droplets is not systematically understood. Boeynaems et al., in their in vitro study use short homotypic RNA (< 50 nucleotides) and PR-repeat peptides to show that RNA can dictate material properties of droplets in an RNA-composition-dependant manner^25^. Maharana et al, demonstrated the ability of RNA to promote (at low RNA:protein ratio) or prevent droplet formation (high RNA:protein ratio)^26^ by prion-like RNA-binding proteins. These studies typically focus on homotypic self-assembly of RNP-protein components in the presence/absence of RNA, and not on multi-component phase separation.

Computational studies in most cases addressed equilibrium aspects of phase separation. Typically, the focus of these studies is to understand how factors such as valency and strength of interactions can tune phase behavior of multivalent biomolecules. Equilibrium studies by Harmon et al. analyzed the physical determinants of phase separation using on-lattice spacer-sticker proteins ^27^. Off-lattice approaches include use of patchy particles to model multivalent proteins, in order to describe how properties of scaffold (large valency proteins) and client (smaller valency) proteins can define phase behavior^28^. In particular, the recent work by Espinosa et al. employs the patchy-particle approach to establish the importance of molecular connectivity in the phase separation of scaffold-client mixtures^29^. A common assumption in existing computational approaches is that the distinguishing feature of scaffolds and clients is their valency alone, and that all adhesive sites can participate in attractive interactions with any other free adhesive site. While a valuable first step in addressing a very complex problem, these studies fail to capture the complex reality of protein-protein interactions networks in the cell where each adhesive site often has a specific binding partner. Also, the primarily focus of previous studies has been on studying the phase-behavior of a mixture of multivalent proteins.^27–29^ Coarse-grained computational models have also been extensively used by Thirumalai and co-workers in their seminal works focusing on the thermodynamics of RNA folding^30–32^. However, the role of physicochemical properties of RNA in modulating phase-separation is less well studied computationally.

In this study, we aim to understand the mechanism by which RNA molecules could promote phase separation of a multi-component protein mixture in a length-dependent manner. Here, we employ a two-pronged approach wherein the interacting protein components not only vary in terms of their valency but also have a defined interaction network, mimicing protein-protein interactions within a known RNP condensate, P-body. We use a hard-sphere patchy particle representation of P-body RNA-binding proteins to study the effect of RNA-length on their phase-behavior at low RNA/protein ratios while modeling the RNA molecules as semi-flexible polymer chains. Using Langevin Dynamics (LD) simulations, we probe the ability of this multi-component protein mixture to form large assemblies at varying protein concentrations and interaction strengths. Further, we specifically focus on the mechanism by which RNA molecules can facilitate droplet formation in the the regime of low protein concentrations and weak interaction strengths by promoting interactions involving low-valency components in the RNP-clusters. Our results suggest that long RNA chains can enable assembly of large clusters even at low concentration of protein scaffolds, and for weak inter-protein interactions.

## Model

The primary focus of this study is to elucidate how RNA molecules can influence the assembly of a multi-component mixture of multi-valent proteins into protein-rich clusters. We study how long RNA molecules (modeled as semiflexible homopolymers) at low RNA:protein ratios influence the assembly of multi-valent proteins (modeled as hard-spheres with interaction patches^29^) into dense clusters using Langevin dynamics simulations. Below, we discuss the protein-protein interaction network modeled in this study and the simulation method in detail.

### Modeling the protein-protein interaction network

Membraneless RNA-protein condensates are made of complex network of RNA-binding proteins and RNA. One such example of an RNA-protein condensate are the P-bodies or processing bodies that are associated with mRNA processing ^6,34^. P bodies are broadly made up of two classes of proteins – the high-valency core proteins (or scaffolds) and low-valency non-core proteins (or clients), ^29,33,34^. Unlike *in vitro* experimental studies with engineered multi-valent proteins, the adhesive sites within P-body scaffolds and clients bind uniquely to a specific interaction site on another component protein ^33,34^.

In Fig. 1A, we represent some components of this interaction network schematically. In our minimal model the P-body contains particles of 16 types modeling its known components such as Dcp2, Edc3, Pat1, Xrn1, Lsm1,Upf1, Dhh1 and RNA^34,35^ (Table 1 and Fig.1). Each of these particles has a finite valency, defined by the total number of adhesive sites (“patches”) on the particle (shown as side beads in the schematic Fig. 1A). Each adhesive site has a unique identity wherein it can only interact with a binding site specific to it on another protein. Therefore, each RNA binding protein that is part of the P-body has a fixed valency with respect to every other protein type. Here, valency refers to the maximum number of adhesive interactions of any particular type that a protein can get involved in. The particle types with large valencies (Dcp2, Edc3 and Pat1 in Fig.1) are refered to as core proteins, while the low-valency components – Pat1, Xrn1, Lsm1 and Upf1 are refered to as non-core proteins. Additionally, each protein component also harbors an RNA-binding domain that can engage in 1 interaction with an RNA molecule.

**Table 1.** Protein-protein interactions in P-body proteins (based on study by Xing el al.^33^)

|  | Protein-protein interactions | Protein-RNA interactions | Total interactions | Core/Non-core |
| --- | --- | --- | --- | --- |
| Dcp2 | Edc3 + Pat1(10 in total), Dhh1, Edc1 | 1 | 13 | Core |
| Edc3 | Dcp2, Edc3, Dhh1, Upf1, Pat1 | 1 | 6 | Core |
| Pat1 | Pat1, Dcp2/Xrn1 (same site), Dhh1, Edc3, Lsm1-7, Upf1 | 1 | 8 | Core |
| Xrn1 | Pat1 | 1 | 2 | Non-Core |
| Lsm1 | Pat1, Lsm1 (2) | 1 | 4 | Core |
| Upf1 | Pat1, Edc3, Upf2 | 1 | 4 | Core |
| Dhh1 | Pat1/Edc3 (same site), Dcp2, (Pop2) | 1 | 4 | Core |
| Pop2 | CCR4, (Dhh1) | 1 | 3 | Non-Core |
| CCR4 | Pop2 | 1 | 2 | Non-Core |
| Not2 |  | 1 | 1 | Non-Core |
| Upf3 | Upf2 | 1 | 2 | Non-Core |
| Upf2 | Upf1, Upf3 | 1 | 3 | Non-Core |
| Hek2 |  | 1 | 1 | Non-Core |
| Sbp1 |  | 1 | 1 | Non-Core |
| Edc1 | Dcp2 | 1 | 2 | Non-Core |

**Table 2.** Important Simulation variables and order parameters

| Notation (or Terminology) | Physical Interpretation | Definition |
| --- | --- | --- |
| $\phi_{prot}$ | Volume fraction of proteins in the simulation box. | $\phi_{prot} = N_{tot} * (4/3 * \pi * R^3) / L^3$,<br>R is the size of the protein monomer,<br>L is the length of the cubic simulation box. This is a dimensionless quantity used to represent protein concentrations. |
| $L_{clus}$ | Size of the single largest cluster | |
| $N_{\text{tot}}$ | Total number of protein monomers in the system. | $N_{\text{tot}} = 1000$ in our patchy-particle simulations |
| $N_{\text{RNA}}$ | Number of RNA chains in simulation | Parameter that is varied in our simulations to vary Protein/RNA ratio. |
| $L_{\text{RNA}}$ | Length of semiflexible RNA chains in LD simulations | Represented in terms of number of Nucleotides |
| $R_g$ | Radius of gyration of RNA | |

**Figure 1:**
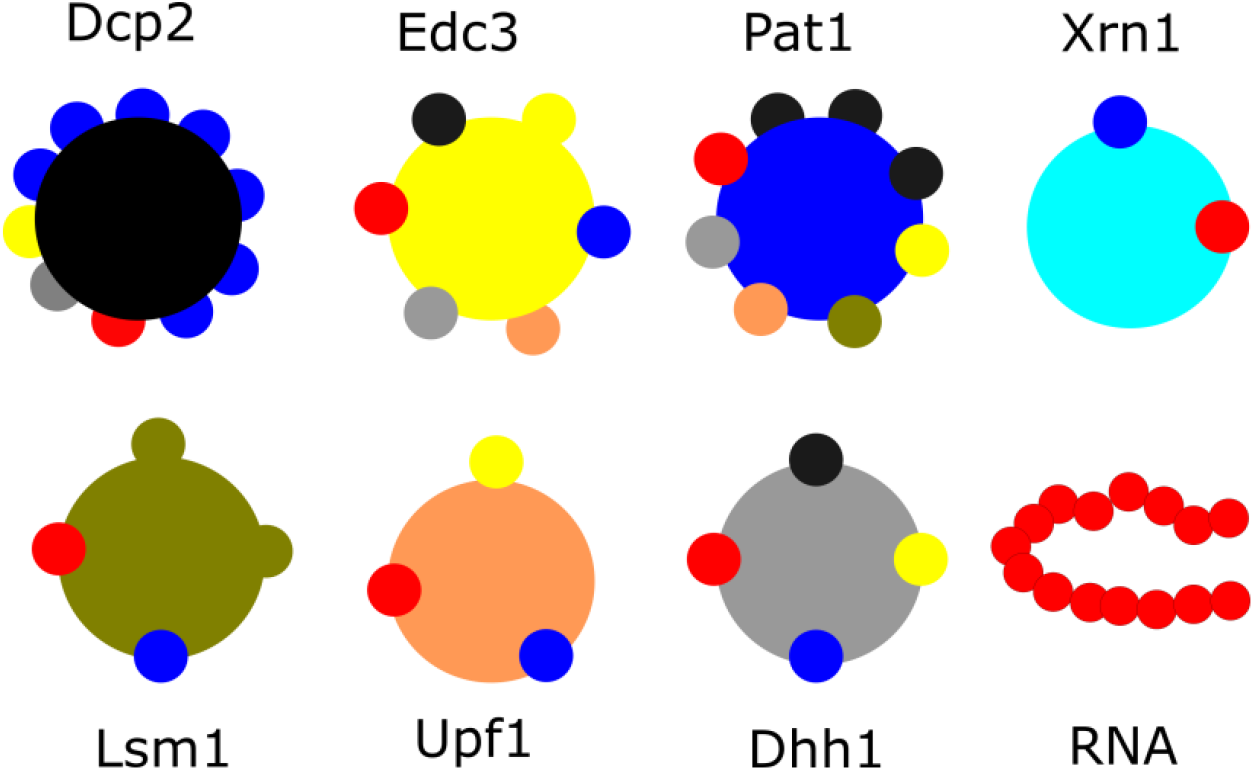
Patchy particle model and the interaction network. Schematic representation of some P-body proteins with different valencies. The colors on the side beads represent an adhesive site that is specific to its complementary adhesive site on another component of the same color. Therefore, each particle has a valency which is specific to every other particle in the simulation. The RNA is modeled as a semi-flexible polymer chain. The length of the RNA chain, L_RNA_, is a parameter that we vary in our simulations.

### Patchy-particle LD Simulations

In order to probe how the RNA-binding proteins assemble into dense clusters in an RNA-dependent manner, we first model these proteins as hard spheres with varying number of adhesive interaction patches on their surface (with interaction preferences defined in Fig.1). Each of these interaction patches can engage in a maximum of 1 adhesive interaction with a complementary patch on another patchy-particle via a continuous square-well potential (see Methods section). The RNA chain, on the other hand is modeled as a semi-flexible homopolymer chain, with each bead of the semi-flexible polymer representing a nucleotide. In our patchy particle LD simulations, we vary the length of the RNA chain, and the RNA:protein ratios to study their role on promoting protein-protein interactions. However, the higher resolution at which the RNA is modeled in the patchy particle LD studies, makes simulations in the limit of large RNA:protein concentrations computationally expensive.

## Methods

### Langevin Dynamics simulations

#### Force Field

The polymer chains in the box are modelled using the following interactions. Adjacent beads are connected by a simple harmonic spring with a spring constant (*k*_*s*_) of 2 kcal/ Å^2^ and an equilibrium bond length (*r*_0_) of 4 Å. The potential energy function describing the connection between adjacent coarse-grained amino acid beads is given by,

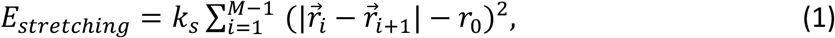

To model electrostatic interactions, we use the standard Debye-Hucklel electrostatic potential with a screening length (*κ*) of 1 nm and a dielectric (*D*) of 80. The Debye-huckel potential has the form,

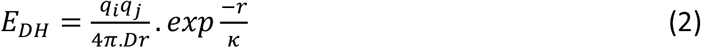

*q*_*i*_ and *q*_*j*_ are the charges of the two interaction RNA s. Each RNA bead carries a single negative charge in our simulations. The RNA chains in our simulation are modeled as semiflexible polymers in which any two neighboring bonds interact via a simple cosine bending potential

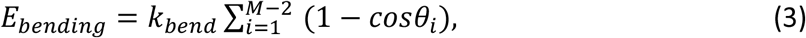

where *θ*_*i*_ describes the angle between *i*^*th*^ and (*i* + 1)^*th*^ bond while *k*_*bend*_ is the energetic cost for bending (bending stiffness of linker).

### Modeling multivalent proteins as patchy-particles

Multi-valency is a key feature of phase-separating proteins, as established by numerous experimental and computational studies. Multi-valent proteins are typically modeled as spacer-sticker polymer chains on- or off-lattice. However, the higher-resolution spacer-sticker polymer chains with higher degrees of freedom are computationally more demanding. However,in a scenario where the microscopic features of the self-assembling chain aren’t central to the problem, multivalent proteins have also been modeled as patchy particles with a defined number of adhesive interaction sites. This MD-Patchy model for multivalent proteins has been extensively used to study phase separation^29^. In this model, the proteins are modeled as hard spheres of diameter *σ*_*HS*_ with a defined number of attractive sites or patches on their surface (Fig. 4A). The central body particle is larger in diameter and is only involved in repulsive hard-sphere interactions. The attractive patches are smaller particles on the surface of this hard sphere, and interact with complementary patches on another particle via an attractive potential that is defined by a continuous square-well potential,

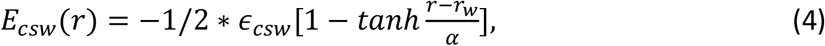

Here *r* is the distance between the centers of two attractive patches and *r*_*w*_ is the radius of the attractive well in the square-well potential. Previous computational studies have established that a cutoff *r*_*w*_ of 0.12*σ* ensures a valency of one per attractive patch^29^. The strength of attraction *ϵ*_*csw*_ is varied in our simulations in the range of 1kT to 5kT. Each multi-valent protein in our Langevin dynamics simulations is, therefore, composed of M+1 particles, where M is the valency of the protein being modeled. It must be noted that these patchy particle proteins are defined as a multi-centre rigid body with no internal dynamics in our simulations. To achieve this, we use the rigid particle definition in LAMMPS^36^. Further, unlike the patchy-particle simulations by Espinosa et al.^29^, each attractive patch in our LD simulations has a specific interaction patch on another protein which is modeled on the complex interaction networks that are typical of RNA-protein condensates such as P-bodies. In other words, not an attractive patch cannot interact with all other attractive patches on other sites. The interaction rules are described in Fig. 1 and Table 1.

### LD Simulations Details

The dynamics of patchy-particle proteins and RNA was simulated using the LAMMPS molecular dynamics package ^36^. In these simulations, the simulator solves for the Newtons’s equations of motion in presence of a viscous drag (modeling effect of the solvent) and a Langevin thermostat (modeling random collisions by solvent particles) ^37^. The simulations were performed under the NVT ensemble, at a a temperature of 310 K. The mass of the multi-valent patchy particle proteins was set to 8000 Da. The size of the rigid, hard-sphere, patchy particles was set to 20Å. The viscous drag in the Langevin dynamics simulations is implemented via the damping coefficient, *γ* = *m*/6*πηa*. Here, m is the mass of an individual bead, ‘*η*’ is the dynamic viscosity of water and ‘a’ is the size of the bead. An integration time step of 15 fs was used in our simulations, and the viscosity of the surrounding medium was set at the viscosity of water (10^−3^ Pa.s).

## Results

### In the absence of RNA, large clusters are disfavored at low volume fractions and weak interactions

Phase-separated condensates such as stress granules and P-bodies are enriched in RNA molecules ^34,35,38,39^. Also, several of the protein components within these structures are known to possess an RNA-binding domain^33^, suggesting that RNA-protein interactions could form the molecular basis of their biogenesis. RNA-binding proteins have also been observed to phase separate at lower threshold concentrations in the presence of RNA ^40–42^. The mechanism by which RNA molecules of varying lengths could influence the assembly of multivalent proteins into condensates, for a defined interaction network, has not been systematically explored. To understand how RNA chains could influence the phase behavior of these complex mixtures of multi-valent proteins in a length-dependent manner, we modeled various P-body proteins ^34^ as hard-spheres with attractive patches^29^ (Fig.1A). The attractive sites on the surface of central core of the protein mimic adhesion sites. The number of adhesive sites on the surface defines the valency of these components. Each adhesive interaction has a specific interaction partner that is defined by the interaction matrix represented schematically in Fig. 1. The RNA molecules, on the other hand are modeled as semi-flexible polymer chains with a negatively charged backbone that prevents homotypic interaction between RNA beads. One adhesive site on each of the 16 patchy-particle protein types is involved in RNA-binding.

We first performed Langevin dynamics simulations of these hard-sphere proteins in the absence of RNA molecules in the simulation box. Two key parameters dictate the phase behavior of these proteins, the total protein volume fraction (ϕ_prot_), and the strength of the protein-protein adhesive interactions (ε_sp_). In our simulations, all multi-valent proteins that make up the multi-component mixture are present at equal concentrations. To model the phenomenon at biologically relevant interaction strengths and bulk protein densities, we vary the interaction strengths (per adhesive interaction), ε_sp_, in the range of 1-5 kT ^27^and protein volume fraction (ϕ_prot_) in the range of 0.01 to 0.1. The ϕ_prot_ values employed in our simulations are of the same order as the estimated optimal protein volume fractions in the cell ^43–45^. In the absence of RNA molecules, at low ϕ_prot_ and weak ε_sp_ (< 4 kT in Fig.2A1), the multicomponent mixture of proteins exists predominantly in the monomeric state or in the form of small clusters (<= 10 monomers). Large clusters (>=100 molecules) are only observed in the limit of strong protein-protein interactions and/or high bulk densities (Fig.2A).

**Figure 2:**
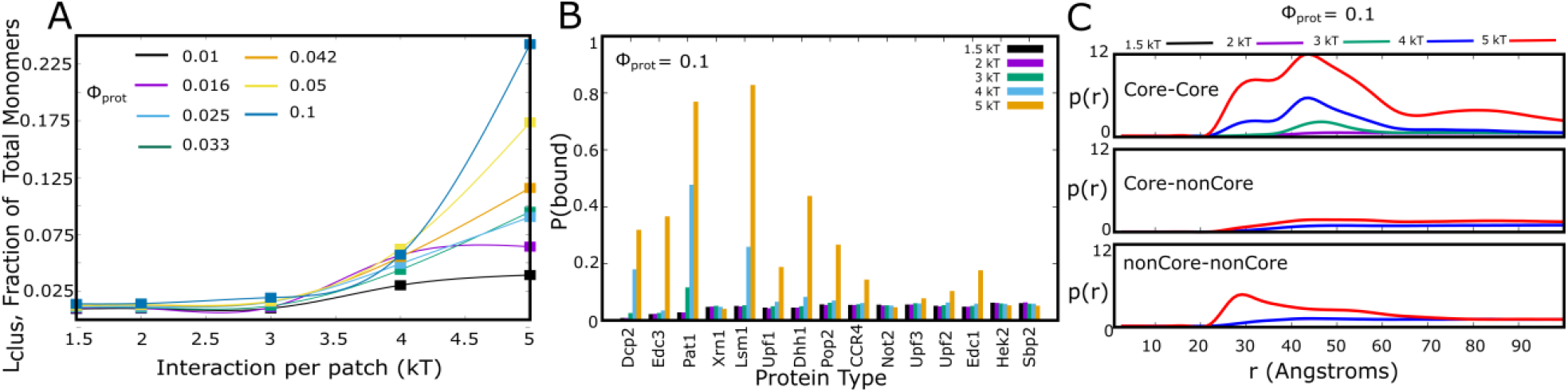
Phase separation in the absence of RNA. A) Assembly sizes in the absence of RNA, for varying protein volume fractions and interaction strengths. The size of the largest cluster (L_clus_) is plotted as a function of interaction strength (per interaction patch). ϕ_prot_ refers to the volume fraction of proteins in the simulation, ϕ_prot_=N_tot_*(4/3*π*R^3^)/L^3^. N_tot_ = 1000, and L is the size of the simulation box. L_clus_ values were computed across 500 different equilibrium snapshots and 5 independent simulation trajectories. B) Probability of finding a multi-valent protein in a cluster that is larger than 10 molecules in size. Dcp2, Edc3, Pat1, Lsm1 are core proteins. Different color bars represent different values of interaction strength, ε_sp_ (per patch). C) Radial distribution function p(r), for core-core, core-non core and noncore-noncore interactions showing the probability of finding two particles of a given type within a cetrain distance of each other (normalized by corresponding value for a pair of ideal gas particles at the same bulk density).

To further quantify the propensity of these multi-valent proteins to assemble into larger clusters, we define a second order parameter, P(bound), which is the probability of finding a protein component in a cluster that is 10 monomers or more in size. In Fig.2B, we plot P(bound) for each multi-valent particle type in the simulation for different values of protein-protein interaction strength. The P(bound) values were computed over several molecules and across 1000 different simulation snapshots. Interestingly, even for a relatively high volume fraction of 0.1, for weaker protein-protein interactions (ε_sp_ < 4kT) all the multi-valent protein components exhibit low values of P(bound). At higher interaction strengths (4 kT and 5 kT in Fig.2B), the two classes of proteins, high-valency core proteins (Dcp2, Edc3, Pat1, Lsm1, Upf1, Dhh1) and the low-valency non-core proteins (Pop2,CCR4,Not2,Upf3,Upf2,Edc1,Hek2), show distinct P(bound) values. The core proteins, at high interaction strengths, are predominantly (>50%) found within large clusters while the non-core proteins are more likely to be observed in the monomeric state (rapid shuttling between the surrounding medium and within clusters). The high bound fraction for the core proteins also gets reflected in the radial distribution function, p(r), for different pairs of particles. The radial distribution function values show the probability of finding two particles of a given type at any distance, normalized by the corresponding values for a pair of ideal gas particles at the same bulk density. As evident from Fig.2C, for high volume fraction, strong protein-protein interactions and in the absence of RNA molecules, the clusters are mediated by core-core interactions. For instance, the orange curve in Fig.2C (ε_sp_ of 5kT) for core-core contacts peaks at twice the value compared to interactions involving non-core contacts.

For strong interactions and large volume fractions, high P(bound) for self-assembled proteins is indicative of minimal exchange of monomers between the protein-rich clusters and the bulk medium in this regime. Functional biomolecular condensates, however, are known for dynamic turnover of components between the droplet phase and the cytoplasm^19,38^. Concentrations of individual proteins within the cell are often in the low nanomolar range^35,46^. Similarly, the protein-protein interactions that govern biomolecular condensation are transient in nature in order to maintain the liquid-like nature of these structures. Therefore, the predominantly core-protein rich clusters observed in the regime of large protein concentration and extremely strong protein-protein interactions might not be relevant to the cellular milieu.

### Long RNA molecules promote clustering of multi-valent proteins even at low concentrations and weak protein-protein interactions

LD simulations in the absence of RNA molecules (Fig.2) reveal that, except for the regime of high protein volume fractions and strong inter-protein interactions, the multi-component system does not undergo phase-separation. Even in this narrow regime of large ϕ_prot_ and ε_sp_, we see core-protein dominant clusters with little or no recruitment of low valency, non-core proteins. This gives rise to some critical questions. Can RNA chains alter the phase behavior of these patchy-particle proteins in the limit of biologically realistic low-to moderate bulk densities of proteins and weak protein-protein interactions? How do non-core proteins get recruited in this regime of low protein concentrations and weak protein-protein interactions? Multivalent proteins that get recruited within biomolecular condensates are often present at low concentrations^21^ in the cell and co-localize RNA chains within them^25,33,47,48^. Therefore, to probe the effect of RNA on the phase-separation of the hard-sphere protein mixture, we performed simulations in the presence of RNA chains of varying lengths – *L*_*RNA*_ of 10, 50 and 120 in Fig.3A. These simulations were performed for an ε_sp_ of 2.5 kT per pairwise interaction patch and a protein volume fraction of ϕ_prot_ = 0.033. For short RNA chains (*L*_*RNA*_ = 10), we do not see an increase in cluster sizes even at large RNA concentrations suggesting negligible phase-separation promoting effect (Fig. 3A, black curve). As we increased the length of the RNA chains from 10 to 50, we see an increase in cluster sizes (L_clus_) for the same number of effective RNA binding sites (Fig. 3A, purple curve). This effect is even more pronounced for *L*_*RNA*_ of 120 (Fig. 3A, green curve), suggesting that longer RNA chains are more effective in promoting the formation of large stable clusters, even at low protein concentrations and weak protein-protein adhesions. In effect, small number of long RNA chains are more effective in promoting self-assembly than large number of short RNA chains at the same number of RNA binding sites.

**Figure 3:**
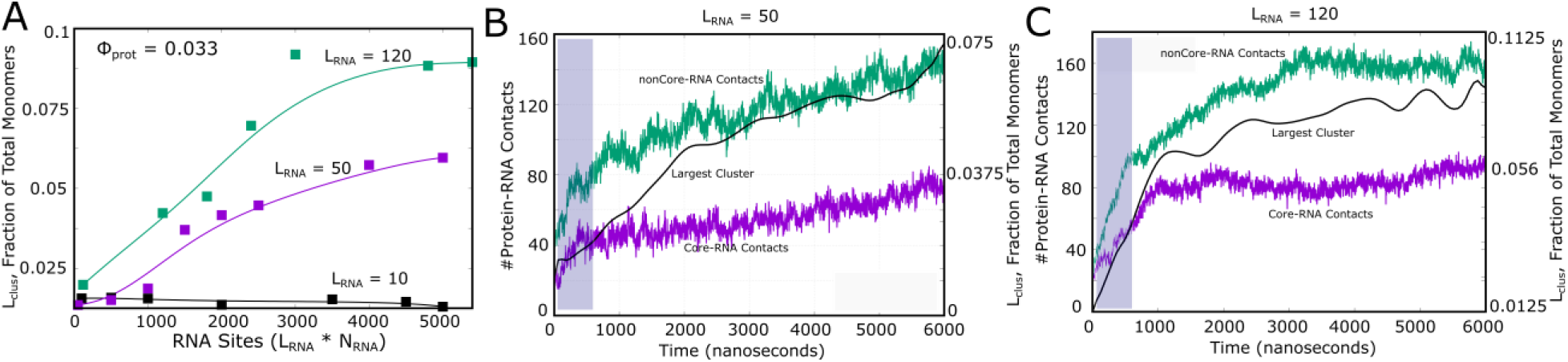
RNA-mediated phase separation. **A)** The values along the x-axis indicate increasing RNA concentrations represented as effective number of RNA beads. Here, *L*_*RNA*_ and *N*_*RNA*_ refere to the length and the total number of RNA chains, respectively. L_clus_ is the size of the single largest cluster in the simulation, at equilibrium, expressed as a fraction of the total number of monomers (N_tot_). The total number of protein particles (N_tot_) in the simulation box was 1000. A L_clus_ of 0.1, therefore, refers to a cluster that is composed of 100 protein monomers. L_clus_ values were computed across 500 different equilibrium snapshots and 5 independent simulation trajectories. Panels **B) and C)** show the RNA-mediated nature of assembly (single trajectory), with an increase non-core RNA-contacts (shaded blue in the figure) preceding the formation of larger sized clusters. ε_sp_ for protein protein interactions was set to 2.5 kT. The interaction strength between any RNA-binding patch and RNA was set to 1 kT (per RNA bead). The curves for the largest cluster size were smoothened using cspline fitting as a guide to eye.

To futher confirm the RNA-mediated nature of self-assembly, we track protein RNA contacts in the system as a function of time, for the core and non-core proteins. As evident from Fig.3B and C, during the early phases of the simulation we observe an increase in RNA-protein contacts involving non-core proteins (green curve). Interestingly, the increase in the size of the largest cluster follows this initial increase in noncore-RNA contacts, suggesting that RNA enables larger clusters by stabilizing non-core proteins within them, even at this low volume fraction (ϕ_prot_= 0.033) and weak interactions (ε_sp_ = 2.5 kT). This effect is more pronounced for long RNA chains (Fig.3C) where we see a sharp increase in RNA-protein contacts followed by a sharp increase in cluster sizes. Overall, these results establish that long RNA chains can mediate droplet formation in a multi-component protein mixture even at low protein concentrations.

### Long RNA chains enable large clusters by recruiting non-core proteins

While the effect of long RNA chains in promoting self-assembly is evident from the large protein-RNA clusters, the mechanism by which RNA chains promote clustering in not well understood. In order to gain further mechanistic insights into the phenomenon, we plot the radial distribution function (p(r)) for different pairs of particles (core-core, core-noncore and noncore-noncore) in our multi-component mixture of RNA and proteins. The p(r) values in Fig.4 refer to the average number of atom pairs of any type (core-core, core-non core and noncore-noncore) found at any radial distance (between r and r+dr), normalized by the equivalent values for a system of ideal gas particles of the same bulk density. A peak in p(r) shows a higher propensity for two particle types to be in vicinity of each other. Sharper peaks are signatures of denser, solid like organization while more difuse peaks indicate liquid-like arrangement. The gpu-based implementation^49^ of the radial distribution function was used to compute the p(r) in Fig.4.

**Figure 4:**
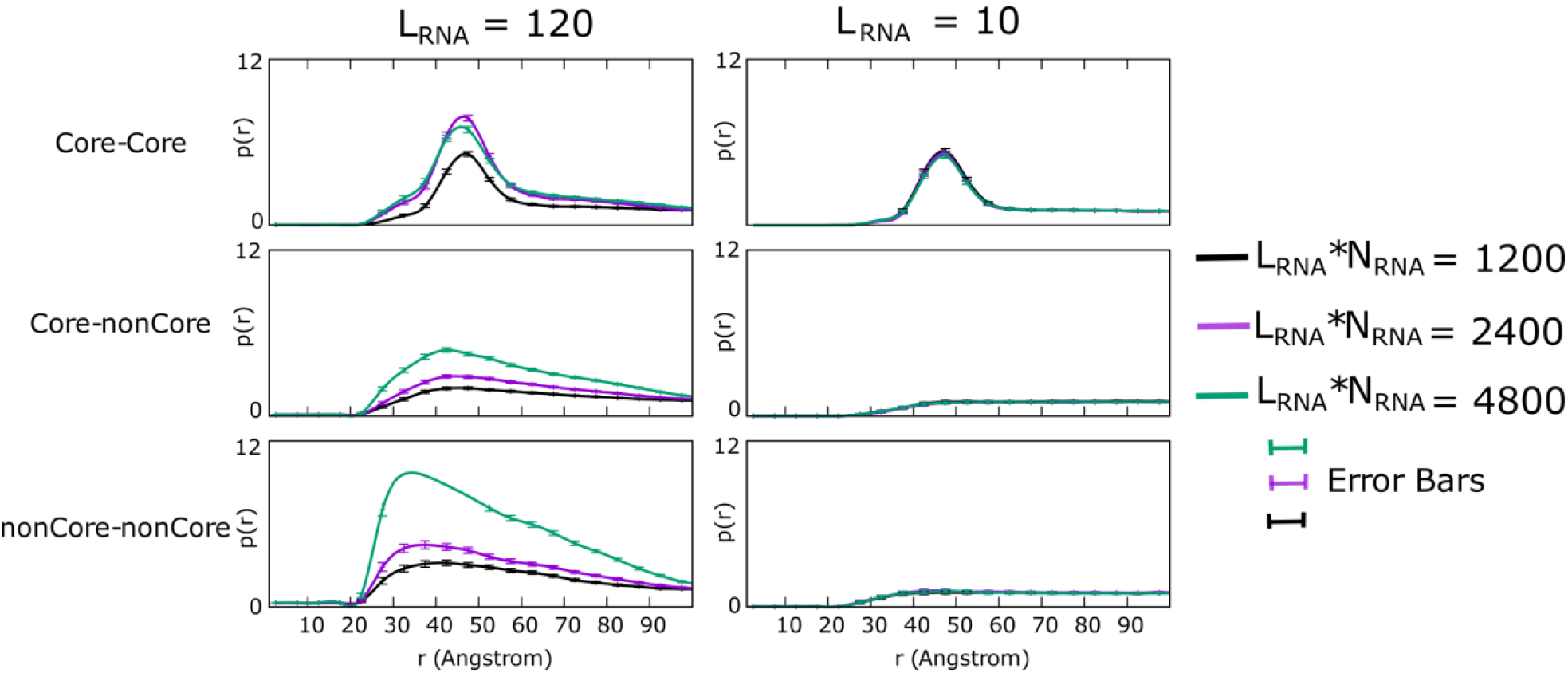
RNA promotes non-core interactions. Radial distribution function, p(r), for core-core (top), core-non core (center) and non core-non core (lowest panel) for L_RNA_ of 10 (right panel) and 120 (left panel). The purple, cyan and black curves represent different numbers of RNA binding sites in the simulation box. The standard errors (S.D/√n) in p(r) were computed over 1000 different equilibrium snapshots.

For low RNA concentrations, the only stable interactions are between core proteins with high valencies (high valency particles in Fig.1A), as evident from the sharp peak in core-core p(r) (black curve for core-core interactions in Fig. 4). In contrast, the p(r) for interactions involving non-core proteins exhibit flat profiles for low RNA numbers (black curve for noncore interactions in Fig. 4). This shows that in the absence of RNA, the clusters can only localize core proteins, and thus are smaller in size (Fig. 3A). As we introduce more RNA chains in the simulation box, the structures that result show a greater likelihood of interactions involving non-core proteins, as evident from p(r) values approaching values of 5-8 (Fig. 4, purple and green curves). For e.g, a p(r) of 5 in Fig.4 indicates that, on an average, a pair of particles is 5 times more likely to be in contact as compared to a system of non-interacting, ideal gas particles for the same bulk density. Further, the longer chain RNA molecules are more effective in promoting non-core interactions as compared to the shorter RNA chains (Fig. 4, left versus right panel). It must, however, be noted that the p(r) profiles for non-core contacts are more diffuse, i.e. liquid-like as compared to the core-core interactions that exhibit a sharper profile (Fig. 4, core-core versus noncore-noncore). Sharp peaks in the radial distribution function are typical of densely packed solids, while liquids exhibit diffuse peaks. Such disparity stems from stronger net interaction strengths for core proteins due to their larger valencies. This finding is consistent with the recent *in vitro* observation of dynamically ‘solid-like’ behavior of large-valency, core proteins which exhibit slower exchange times^33^. Therefore, the p(r) profiles suggest that the presence of long RNA chains promotes larger clusters by facilitating non-core interactions.

Further, we computed the probability, P(bound), of finding individual particle types within large clusters (>10 monomers in size). For short RNA chains (Fig.5A), we observe little or no self-assembly, with P(bound) values not exceeding 0.1 for the component proteins in the mixture. These P(bound) values, in presence of short RNA, are akin to those observed in the absence of RNA, at weak protein-protein interaction strengths (<4kT in Fig.2B). However, as we increase the length of the RNA chains in the simulation box, we see a dramatic increase in P(bound) values across different protein components (Fig.5B and C). Also, unlike the assemblies observed at large volume fractions and strong inter-protein interactions (in absence of RNA, P(bound) à 1), the proteins in RNA mediated clusters exhibit low P(bound) values suggesting rapid exchange between the monomeric and self-assembled states. Overall these results suggest that RNA molecules can act as super-scaffolds that promote phase-separation of multi-valent proteins, interacting via defined networks, in a length-dependent manner.

**Figure.5:**
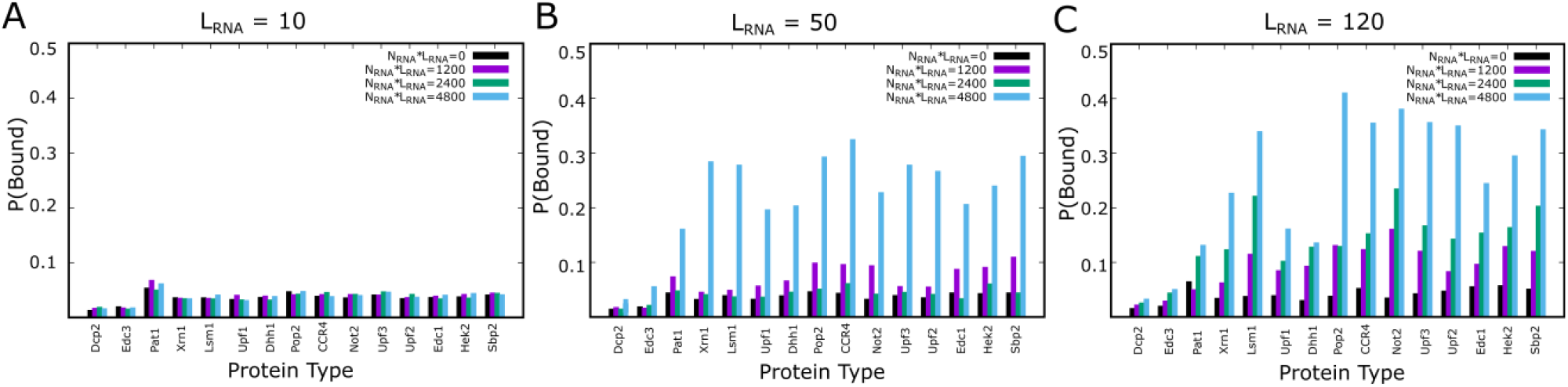
Probability of finding each protein component within a multimeric cluster (>=10 monomers) in presence of increasing RNA concentrations. The three panels A), B) and C) show results in presence of RNA chains of length 10, 50 and 120, respectively. Higher values of P(bound) indicate a greater likelihood of a component being found within RNA-protein clusters.

### RNA-mediated clustering of proteins causes compaction of RNA

One of the key functions of RNA-protein condensates is the sequestration of mRNA molecules stalled in translation (stress granules^50^) or the processing of RNA^34^. Single molecule experimental studies have revealed significant conformational changes in mRNA molecules depending on its translational state and cellular localization. To further test whether the coarse-grained RNA chains in our simulations exhibit behavior that is consistent with experiments, we probe the conformational dynamics of RNA chains in our simulations, for increasing volume fraction of multi-valent proteins. In Fig.6A, we show the distribution of the radius of gyration of RNA chains in the simulation at equilibrium, computed over 1000 different equilibrium snapshots and multiple RNA chains in the simulation box. As we increase the volume fraction of proteins within the simulation box from 0.01 to 0.1, the distribution of RNA sizes shifts to lower values, with the peak shifting from 40 to 20 A, as protein volume fraction approaches 0.05 or higher. In Fig.6B, this gets reflected in the sharp decline in <Rg> at higher volume fractions, suggesting that RNP-assembly results in compaction of RNA, facilitated by RNA-protein contacts within protein-rich clusters. Interestingly, this is consistent with experimental studies which show that translationally inhibited mRNA molecules get sequestered within stress granules in a compact, circularized configuration^50^.

**Figure 6:**
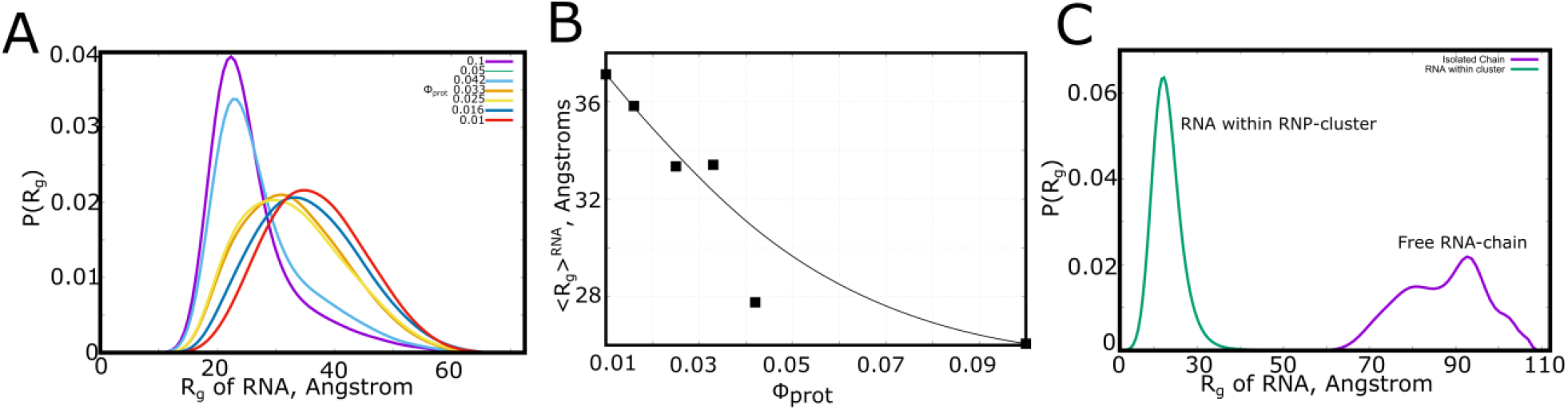
RNA compaction within cluster. A) Distribution of radius of gyration of RNA chain at increasing bulk density of proteins. At higher volume fractions, where RNA-protein clusters form, the RNA chains assume a more compact configuration. B) Comparison of conformational statistics (radius of gyration) of a sequestered RNA chain within an RNP-cluster and a free RNA-chain. Statistics in this figure were computed over 1000 different equilibrium snapshots and multiple RNA chains in the simulation box.

**Figure 7:**
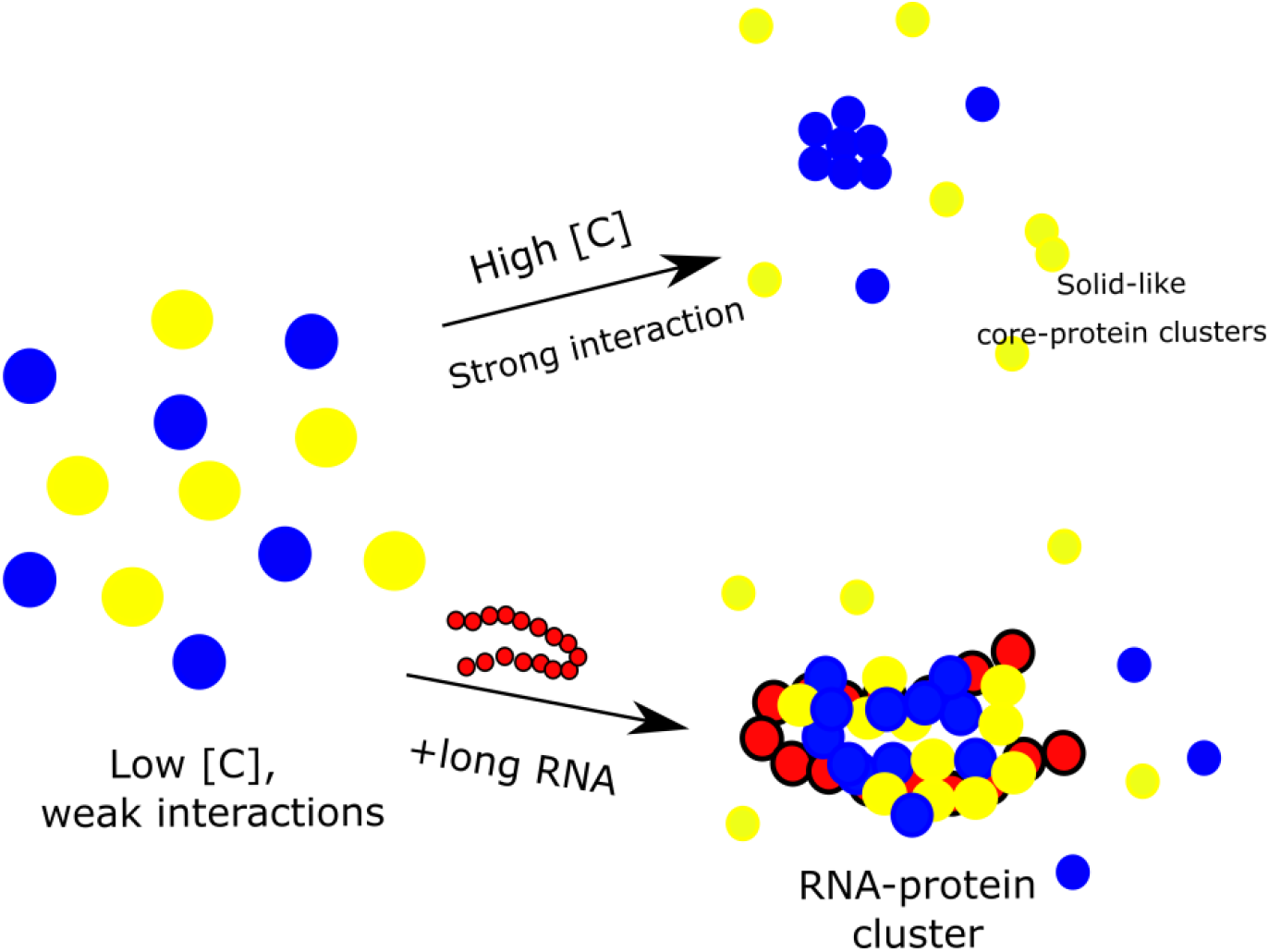
Summary. Addition of long chain RNA molecules can promote large protein-rich clusters even at low concentrations [C] and weak protein-protein interactions. These clusters enrich non-core proteins (yellow balls) more than the clusters that are observed at large protein concentrations and strong protein-protein interactions. Blue balls represent core proteins.

## Discussion

In vitro and in silico studies have established the ability of several RNP condensate components to form droplets even in the absence of other condensates components ^41,51^. An important caveat from these studies is that the phase separation of RNA-binding proteins in vitro and in vivo typically occurs at concentrations that are typically higher than their normal endogenous cellular concentrations^19–21^. The complex nature of the intracellular space and the multi-component nature of the condensates necessitates a systematic exploration of non-protein components in driving the phase-separation of protein-RNA clusters. RNA molecules promote phase-separation of several RNP-proteins at low-moderate RNA:Protein ratios, and are also enriched within condensates like stress-granules and P-bodies^34^. Yet, these model studies are often far from the cellular reality where the phase-separated state is a complex mixture of multi-valent proteins, each with a different valency and defined adhesive interactions ^33,34^.

We employ a coarse-grained computational approach that allows a systematic bottom up exploration of RNA-mediated assembly of RNA-protein clusters for a broad range of conditions (protein volume fraction, protein-protein interaction strength, RNA-length). This approach is similar in philosophy to the in vitro study by Banani et al ^17^ where they design scaffold and client peptides based on differing repeat lengths. However, the scaffolds and clients in the Banani model only differ in valencies but do not incorporate any of the complex interaction networks that are typical of protein-protein interactions within condensates. Our phenomenological simulations are, therefore, an attempt to extend the understanding of a client-scaffold system to a more complex reality with defined interaction networks (Fig. 1) to study the phase transitions in a multi-component mixture. Espinosa et al have previously used a similar coarse-grained computational approach to study the phase behavior of a multi-component mixtures of scaffold and client proteins^29^ in response to critical parameters such as pH, salt concentration, temperature etc. However, in a variation from the Espinosa study, the scaffold and client protein particles in our phenomenological model do not just vary in valency but also in the identities of the adhesive sites. Each adhesive site on the interacting particles in our patchy-particle LD has a unique binding partner on another protein --a coarse-grained representation of the network of specific interactions within the P-body.

A systematic exploration of the behavior of this multi-component system for a range of protein volume fractions and protein-protein interaction strength reveals that, in the absence of RNA molecules, clustering of proteins is observed only at large volume fractions and strong protein-protein interactions. The protein-rich clusters that assemble in this regime of concentrations and interaction strengths are predominantly enriched in core proteins (Fig.2B, ε_sp_> 3kT). Strikingly, the regime that favors large clusters in the absence of RNA features long dwelling times of core-proteins within these clusters. Large valency proteins (for e.g Pat1 in Fig.2B), in this concentration regime are almost entirely localized within these clusters, and exhibit negligible turnover (Fig.2B, P(bound) à 1 for ε_sp_= 4kT and 5kT). The non-core proteins, on the other hand, do not participate in these clusters (Fig.2B, P(bound) << 1). This raises an interesting paradox. Individual phase separating proteins are found at low concentrations in the cell. Moreover, intracellular droplets are liquid-like and exhibit fast turnover dynamics, suggesting weak, promiscuous interactions^52^ stabilize these assemblies. How do cells achieve phase-separation in the endogenous low-concentration and weak interaction regime? Earlier studies showed that mRNA molecules lower the threshold concentrations for phase separation (at low RNA:protein ratios) of prion-like RNA-binding proteins in vitro^26^. Further, threshold concentrations for homotypic phase separation of proteins in vitro can be several fold higher than their cellular concentrations^21^. In vitro reconstituted droplets are also prone to exhibit maturation into solid-like assemblies compared to the more dynamic in vivo counterparts that assemble at lower concentrations^40^.

We systematically probed whether RNA can promote phase-separation of multi-component protein mixture in the regime of low protein volume fraction and weak protein-protein interactions. Consistent with model *in vitro* studies, the introduction of long-RNA molecules facilitates assembly of the multi-component protein mixture to into large clusters of 10 particles and more even at a low protein concentration and weak protein-protein interactions. A detailed exploration of the process of assembly reveals that RNA facilitates larger clusters in this regime by stabilizing interactions involving non-core proteins, as evident from Fig.3B and C where noncore-RNA contacts precede the formation of large clusters! This is also consistent with the radial distribution functions which show an increased likelihood of interactions involving non-core proteins in the presence of RNA. Interestingly, the enrichment fraction P(bound), shows a significant increase for all protein components. However, unlike clusters observed in the absence of RNA in the regime of large protein concentrations and strong interaction, the the P(bound) values do not approach 1 for RNA-dependent clusters. This indicates that the clusters formed in the presence of RNA chains exhibit greater turnover of individual components. In vitro phase separation in the absence of RNA can therefore result in droplets with altered composition as well as intra-droplet dynamics compared to their in vivo counterparts.

The RNA-mediated demixing exhibits a strong dependence on RNA chain length, with long RNA chains (Fig.3A) enabling larger clusters even at low protein volume fractions. These results suggest that long RNA-molecules behave like high-valency super-scaffolds which can act as a hub that networks core and non-core proteins. This behavior of long RNA chains is analogous to the condensate-stabilizing effect of multi-valent spacer-sticker ligands in a recent study by Ruff *et al*^53^. Similarly, the patchy particle simulation study by Espinosa et al suggests that high valency, promiscuous binders can efficiently promote phase separation^29^. These studies, however, model scaffolds and client molecules as collapsed structures (patchy hard spheres)^29,53^. In the protein-protein interaction network modeled in our current work, the large valency proteins (core proteins in Table 1) cannot promote large clusters in the low endogenous concentration regime because of their specificiy. RNA chains, on the other hand, due to their ability to bind to several proteins, can act as physical crosslinks between proteins in the cluster, allowing them to interact with their partners in this high-density phase. Further, the weaker strength of RNA-protein interactions (1kT) compared to protein-protein interactions can result in dynamic forming and breaking of interactions within the condensate, thereby providing a liquid-like environment. The structures that assemble in the absence of RNA (strong inter-protein interactions) are also different in composition to RNA-mediated assemblies for weak protein-protein interactions. In their seminal study on stress granule RNA-enrichment ^22,24^, Khong et al show that RNA enrichment within condensates is strongly correlated to RNA-length, with long RNA chains (> 1 kB) more likely to be enriched within SGs. Formation of membraneless organelles is often a response to stress, and dynamic changes in RNA metabolism can trigger the reversible assembly of these structures^54^. Spatiotemporal regulation of RNA concentration, and selective enrichment of mRNA in a length-dependent fashion, therefore, make RNA molecules efficient enhancers and modulators of reversible phase separation in the regime of low protein concentrations and promiscuous interactions.

## Acknowledgements

The authors would like to thank Ranjith Padinhateeri and Mark Miller for useful discussions. This work is suppoerted by NIH GM 068670

## TOC Figure

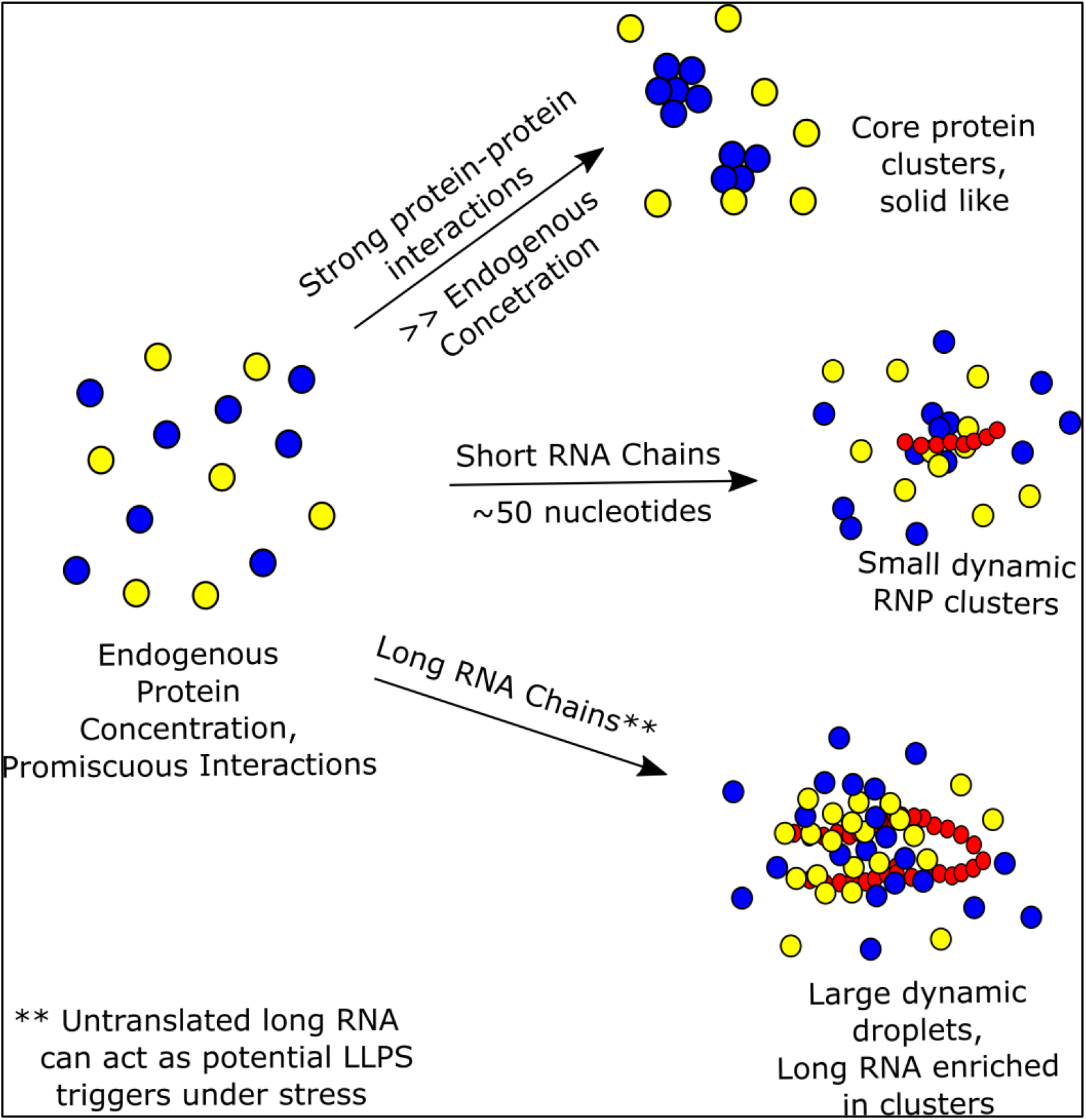

## Notes

### Competing Interest Statement

The authors have declared no competing interest.

